# The critical roaming hypothesis: arousal-driven transitions across critical lines reproduce human functional connectivity dynamics

**DOI:** 10.64898/2025.12.29.696846

**Authors:** Anagh Pathak, Demian Battaglia

## Abstract

Ongoing brain activity displays rich temporal variability associated with efficient cognition, with functional connectivity (FC) continually reconfiguring over time. The resulting functional connectivity dynamics (FCD) specifically show complex, fat-tailed statistics that alternate between persistent epochs and faster reconfiguration transients. While nonlinear whole-brain models tuned nearby a critical point have reproduced some aspects of FCD, they fall short of capturing its full temporal complexity. We propose that slow fluctuations in arousal offer a biologically plausible mechanism for exploring critical regimes in large-scale brain dynamics and thus enrich FCD. Using a connectome-based model of coupled cortical populations, we identified phase boundaries where system dynamics transition between regimes of faster or slower FCD. We then phenomenologically incorporated arousal changes, modeling them as stochastic fluctuations in key parameters such as cortical excitability, input gain, and noise amplitude. This non-autonomous formulation enables the system to roam dynamically across regime boundaries, flexibly tuning its distance from critical transition lines and producing intermittent transitions that mirror the stochastic evolution observed in empirical FCD. Fitting these models to human resting-state fMRI and performing model comparison, we find that arousal-driven models more accurately reproduce the distinctive quantitative features of FCD with the greatest improvements coming from the previously poorly accounted fat-tailed portions of the distributions. Together, these results suggest that arousal fluctuations –likely mediated by changes in neuromodulatory tone – shape the brain’s attractor landscape over time, expanding the repertoire of accessible functional network states and providing a mechanistic basis for the complexity of spontaneous functional dynamics.

## Introduction

Even during quiet wakefulness, the mind is rarely still—its ongoing mentation drifts and alights like a bird in flight^1^. The neural counterpart of this restless stream of consciousness is echoed in Functional Connectivity (FC)–the pattern of correlated activity across distributed brain regions measured with fMRI–which itself fluctuates dynamically over time^2^. Much like the complex trajectories of real bird flight^3^, the temporal evolution of functional connectivity (FC) can be conceptualized as a random walk through a high-dimensional connectivity space, where each step reflects the reconfiguration of large-scale network interactions^4,5,6^. Empirical analyses have shown that the distribution of FC step lengths deviates substantially from a Gaussian form, exhibiting fat-tailed statistics that signal complexity^4^. These dynamics alternate between “knots” and “leaps”, i.e. epochs of transient FC stabilization and rapid reconfiguration, respectively^4^. Importantly, such temporal complexity in functional connectivity dynamics (FCD) correlates with individual differences in aging and cognition^7,8,4,5^ and can serve as a neuromarker of cognitive decline in neurodegenerative diseases^9,10,11,12,13^.

To understand the origin of these statistical features, one must consider the underlying dynamical system that generates them. A helpful intermediate-level description is provided by the metaphor of an *attractor landscape*, in which brain activity evolves like a particle moving across a manifold whose geometry constrains its trajectory^14^. Within this framework, the fat-tailed distribution of FCD step lengths can arise naturally from multi- or metastable dynamics–such as heteroclinic cycles producing slow drifts punctuated by rapid transitions, or noise-enabled switching among stable FC configurations ^15,16,17^. Such behaviors can emerge from the autonomous collective dynamics of coupled, stochastic, nonlinear neural populations, which constitute a dominant modeling approach to explain the temporal structure of FCD ^15,17^. In this paradigm, structured FCD reflects operation at a single, fixed Dynamical Working Point (DWP), hypothesized to lie near a critical point of the system’s dynamics^18,19,20^.

A complementary and less explored mechanism involves changes in *arousal*. Here, we use arousal to denote a global brain state that shapes vigilance and behavioral responsiveness through diffuse neuromodulatory influences on cortical activity and input integration. Multiple systems contribute to arousal^21,22,23^, including cholinergic and noradrenergic pathways that project broadly across the cortex and can rapidly reshape the attractor landscape of neural dynamics^24^. Computational models linking neuromodulatory effects to biophysical parameters show that activation of these systems can shift the brain between more integrative or segregative network states^25^. Such transitions–potentially sharp even in response to smooth fluctuations in neuromodulatory tone–are facilitated by critical boundaries separating distinct dynamical regimes, with neuromodulatory inputs acting as control signals that steer the system’s DWP through parameter space. The primary control knobs of this process—parameters such as cortical excitability or neural gain—represent plausible sites of neuromodulatory action, enabling the brain to expand or contract its repertoire of accessible network configurations in a task-contingent manner, traversing critical transition lines with qualitative changes in dynamics.

In time-resolved FC studies, arousal is often treated as a nuisance factor; yet, unlike motion or physiological noise, arousal fluctuations have a genuine neural origin^26^. Recent animal work shows that arousal can be captured by a low-dimensional variable–such as pupil diameter–that explains a substantial fraction of ongoing, large-scale neural activity^27^. While large swings in arousal accompany sleep–wake transitions, more subtle modulations also occur during quiet wakefulness, suggesting that continuous arousal fluctuations may act as a latent driver of spontaneous network reorganization^28,29^. Extending the attractor metaphor, brain activity would no longer unfold on a static landscape; rather, the landscape itself would be continually reshaped as the system’s DWP drifts under the influence of fluctuating arousal, altering its instantaneous distance from the critical transition lines that structure the system’s collective behavior ^30,31,32,33^.

Here, we adopt a whole-brain computational modeling approach to assess whether dynamic complexity at a DWP near criticality is sufficient to generate structured and fluid FCD, or whether temporal fluctuations of the DWP are also required to account for the empirically observed organization of FCD. Specifically, we simultaneously fit multiple quantitative features of human resting-state fMRI FCD using two alternative families of connectome-based whole-brain models: first, autonomous dynamical systems in which activity unfolds on a fixed landscape; and second, non-autonomous, arousal-driven models in which the dynamical landscape itself changes over time.

We focus in particular on how well each model reproduces the fat tails of the empirical FCD fluidity and speed distributions, rather than just their mean values. Although both model classes capture general features of FCD, only the time-varying (non-autonomous) models naturally generate the very slow-speed events and the “viscous” reconfiguration patterns observed in empirical data—phenomena reflected in frustrated link-to-link interactions along FCD trajectories^34,11,12^. This is notable because precisely these reductions in dynamical fluidity—robust across multiple approaches yet poorly captured by previous modeling frameworks—have been proposed as markers of cognitive decline in aging, demanding task states, and neurodegenerative disease^4,5,6,11,12,13^.

Our findings therefore suggest that intrinsic FCD is sculpted by ongoing fluctuations in neuromodulatory tone associated with arousal. They also refine earlier theories proposing that the brain sits near a single critical boundary. Instead, our fitted simulations indicate that the dynamic working point roams across a broad inter-critical zone— spanning a large expanse along the ignition threshold^35^—giving rise to non-monotonic changes in FCD fluidity as arousal varies.

## Results

Unlike previous approaches that segment functional connectivity dynamics (FCD) into discrete transitions between quasi-stable FC states ^36^, we describe FCD as a smooth, continuous flow through a space of continually morphing connectivity configurations ^4^. Conventional analyses of static FC emphasize the spatial structure of connectivity networks while discarding most temporal information. An alternative perspective collapses each FC matrix into a single point in the high-dimensional space of all possible FC configurations—connectivity space. The erratic motion of this point through time can then be viewed as a stochastic trajectory or random walk across the manifold of possible network states. Most prior modeling studies have focused on reproducing the distribution of FCD values, while neglecting the sequential dependencies that define the temporal organization of FC reconfigurations^37,38^. Randomly shuffling the FCD streams, for instance, preserves the overall distribution but destroys its temporal structure—thus erasing the random-walk nature of the process ^4^. To capture this essential aspect of FCD, we extend model fitting beyond the FCD distribution itself to include its step-length distribution, quantified as *FCD speed* (Fig1). This metric characterizes how rapidly the system traverses connectivity space and the tails of this distribution encode rare but dynamically important transitions (corresponding to FCD “knots” and “leaps” ^4^).

In the attempt to reproduce a rich FCD structure *in silico*, we first focus on a type of mean-field model (MFM) that has been extensively applied to characterize large-scale brain dynamics and reproduce key features of functional connectivity^39,15,38,40^. The model contains several parameters that can plausibly serve as control knobs for arousal, including cortical excitability and neural gain (which together set responsiveness and the relationship between synaptic input and population firing rate), the amplitude of stochastic noise, and the strength of local recurrent coupling within each neural population. Owing to a diversity of model parameters and nonlinear architecture, the MFM provides multiple avenues through which neuromodulatory input can transform system dynamics, making it a suitable platform for probing how arousal reshapes large-scale functional organization. In the following, we refer to the autonomous formulation as the mean-field model (MFM) and to its time-dependent extensions as tMFM, where *t* denotes the explicit temporal modulation introduced by arousal. Further, we label each tMFM variant according to the specific parameter endowed with temporal dependence: tMFM*_G_* for the global scaler of inter-regional coupling (*G*, coupling gain), tMFM*_n_* for background noise amplitude (*σ*), tMFM*_a_* for the gain parameter of regional sigmoidal response functions (*a*), and tMFM*_w_* for the intra-regional recurrent connectivity strength (*w*).

### Existence of critical transitions in FCD fluidity

Neuromodulatory systems are thought to dynamically tune the brain’s operating regime, enabling flexible transitions between distinct functional states ^25,41^. The parameters of the mean-field model (MFM) represent potential targets of such modulatory control. Given the non-linear structure of the MFM, we further hypothesized that variations in key parameters could drive the system across critical boundaries, thereby reshaping the dynamic landscape of large-scale activity. To examine this, we asked how individual model parameters influence overall network dynamics and, in particular, the speed of fluctuations in functional connectivity dynamics (FCD).

We systematically varied six model parameters—global coupling strength (*G*, also referred to as global excitability following^25^), noise amplitude (*σ*), local recurrent gain (*w*), and the three coefficients (*a*, *b*, *d*) governing the sigmoidal input–output function of each neural population (slope, threshold and smoothness). Among these, *G* and *σ* represent global parameters controlling inter-regional coupling and stochastic drive, whereas *w*, *a*, *b*, and *d* define local circuit properties shared across regions^39^.

The parameter sweep revealed that *G* and *σ* exert the strongest influence on FCD speed, giving rise to distinct dynamical regimes (Fig. 2A). In the *G*–*σ* plane, a wedge-shaped region emerged where FCD speed —here tracked by the obtained median value *ν*_50_— was markedly reduced, suggesting the presence of a critical transition zone. Notably, entry into this regime required a finite level of stochastic input, consistent with a form of stochastic resonance^42^. The low-speed wedge coincided with elevated temporal rate variability (mean standard deviation of firing rates; Fig. 2C), whereas its lower boundary aligned with a rate instability that produced increased spatial rate variability across regions (Fig. 2B). Our results are consistent with previous parametric studies of the Wong–Wang whole-brain model, which identified two rate boundaries at higher global coupling: a lower threshold is associated with the critical *ignition* line (*G*_−_), at which some regions, due to their dense neighborhood, are able to self-sustain themselves in a “up” state with large firing rate, even in the absence of strong external drive ; and an upper *flaring* threshold marking a rate instability (*G*_+_) beyond which all regions saturate in their high firing rate regime ^15,35^. These critical transition boundaries were here more formally identified using an unsupervised clustering procedure (Supplementary Section 4).

Across all examined parameters, the non-linear nature of the MFM generated sharp phase transition boundaries, underscoring its sensitivity to small perturbations in parameter state (Fig 2E-L). These findings indicate that modest parameter changes can reorganize the dynamical regime of large-scale brain networks.

### Arousal as a global modulatory drive in the mean-field model

Could fluctuations in arousal constitute the biological mechanism driving small parameter changes that traverse critical regimes and thereby reorganize large-scale dynamics? We hypothesized that slow, endogenous variations in arousal modulate key model parameters such as global coupling (*G*) and noise amplitude (*σ*)—thereby steering the system through different regions of the parameter space identified in the prior sweep.

To test this, we introduced an explicit arousal variable evolving as a smooth, time-dependent signal that parametrically influenced model parameters. For generality, we modeled this arousal signal as a truncated Ornstein-Uhlenbeck process, defined by a baseline value, mean-reversion rate, and stochastic drive. The baseline tone reflects empirical observations from pupil diameter and locus coeruleus firing, which indicate distinct tonic and phasic neuromodulatory modes ^43^.

This formulation transforms the mean-field model from autonomous to non-autonomous—i.e., from static to time-dependent—, allowing arousal to act as a global control parameter that continuously modifies the system’s DWP and thereby, its dynamical landscape. We then assessed whether arousal-driven modulation reproduces the temporal structure and variability of functional connectivity dynamics (FCD) observed in human fMRI data, and how it compares to the autonomous model. To probe which portions of the FCD and FCD-speed distributions are most influenced by neuromodulatory control, we parametrized FCD and FCD-speed distributions using percentiles (5*^th^*, 25*^th^*, 50*^th^*, 75*^th^*, 95*^th^*) and used these as fitting objectives in a genetic algorithm^44^ (see *Methods*).

Comparing autonomous and non-autonomous models across 200 recordings (see details for five representative subjects in Fig. 3B–C and summary statistics in Fig. 3D), we found that incorporating arousal-driven modulations of model parameters significantly improves model fit, even when penalising for the additional parameters introduced by the Ornstein–Uhlenbeck process. Focusing on the tMFM*_G_* model (in which *G* is under neuromodulatory control), most of this improvement stems from its ability to accurately capture the tails of the FCD fluidity *χ* and speed *ν* distributions —as illustrated by the example distributions shown at the bottom of Fig. 3B— and in particular the left tail (5th percentile). This finding is consistent with our hypothesis that neuromodulatory inputs selectively enhance slow dynamical regimes (Fig. 1). At the same time, we observed that the tMFM*_G_* model also markedly improves the representation of the right tail of the FCD-speed distribution (95th percentile of *ν*), suggesting the occurrence of abrupt crossings of critical transition boundaries that are absent in the autonomous model (Fig. 3C,D).

**Figure 1:**
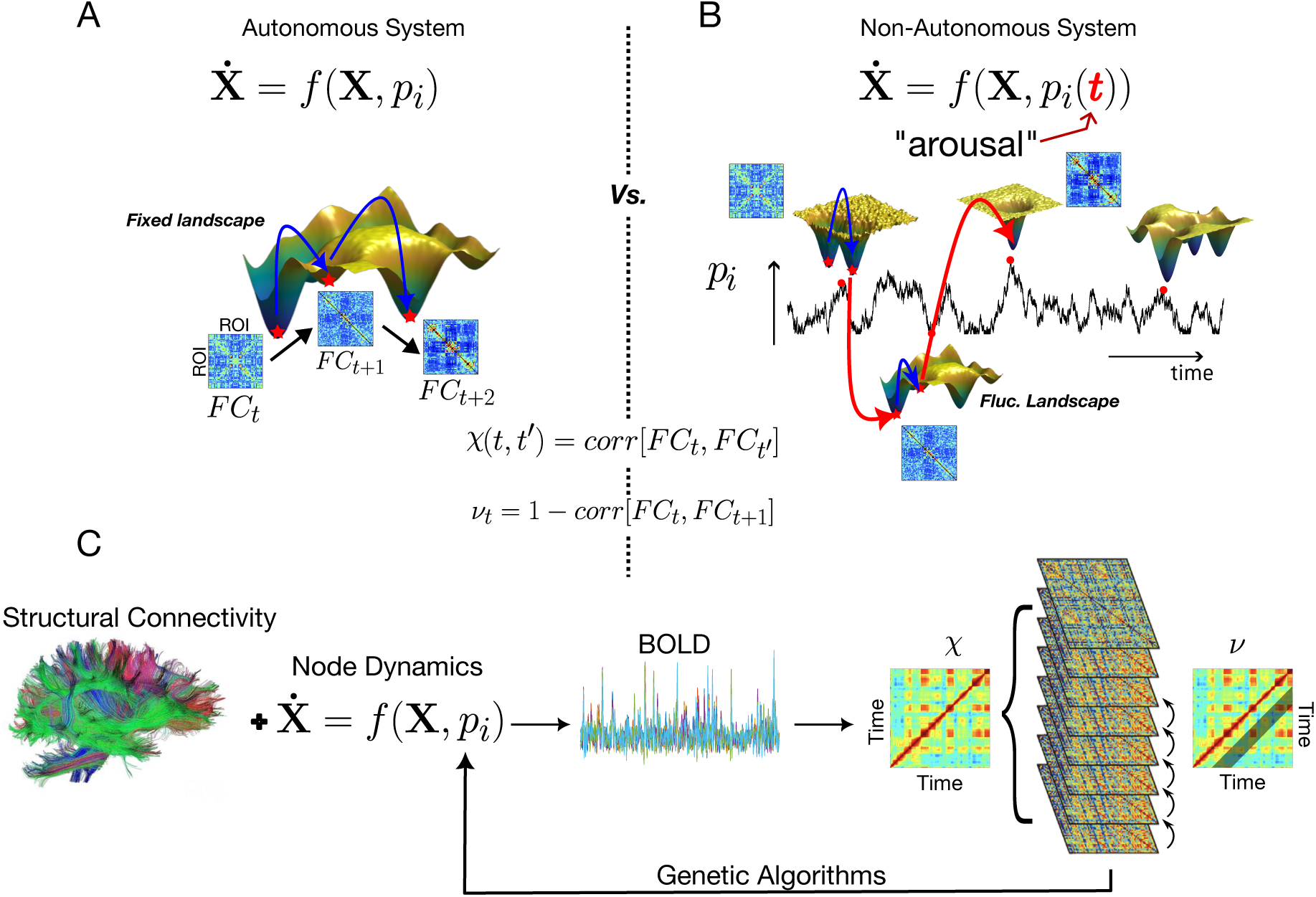
Calibrating Autonomous vs. Time-Varying Models to FCD Statistics: We compare the performance two classes of non-linear models at capturing the temporal statistics of on-going FCD as reflected in the distribution of FCD matrix entries (*χ*) and of sequential FCD variations, or FCD speed (*ν*). A) The activity of each brain region is described by autonomous ordinary differential equations (mean field model); each brain region receives inputs from other brain regions weighted by connection strength (as specified by diffusion imaging). The collective dynamics of this network gives rise to FCD, whose variabiity is tracked by the entries (*χ*) of a recurrence matrix which gives the correlation between the windowed FCs across time frames. FCD speed (*ν*) represents then the correlation distance between successive FC frames. The two metrics *χ* and *nu* thus together overall FCD variability and its temporally ordered flow. B) To mimic the influence of arousal fluctuations, we use the same set of equations as in panel (A) but make some of the coefficients of the model time-dependent (fluctuating as a Ornstein-Uhlenbeck process). C) Using a Genetic Algorithm (Materials and Methods), the model was calibrated to a subset of resting-state fMRI recordings from the Human Connectome Project. The mean-field model’s internal parameters were optimized to align simulated dynamics with empirical observations, using the 5*^th^,* 25*^th^,* 50*^th^,* 75*^th^,* 95*^th^* percentiles of *χ* and *ν* as fitting targets, to account simultaneously for typical as well as extreme transient behavior. For the non-autonomous (time-dependent) model, the parameters governing the Ornstein–Uhlenbeck process were additionally fitted.

**Figure 2:**
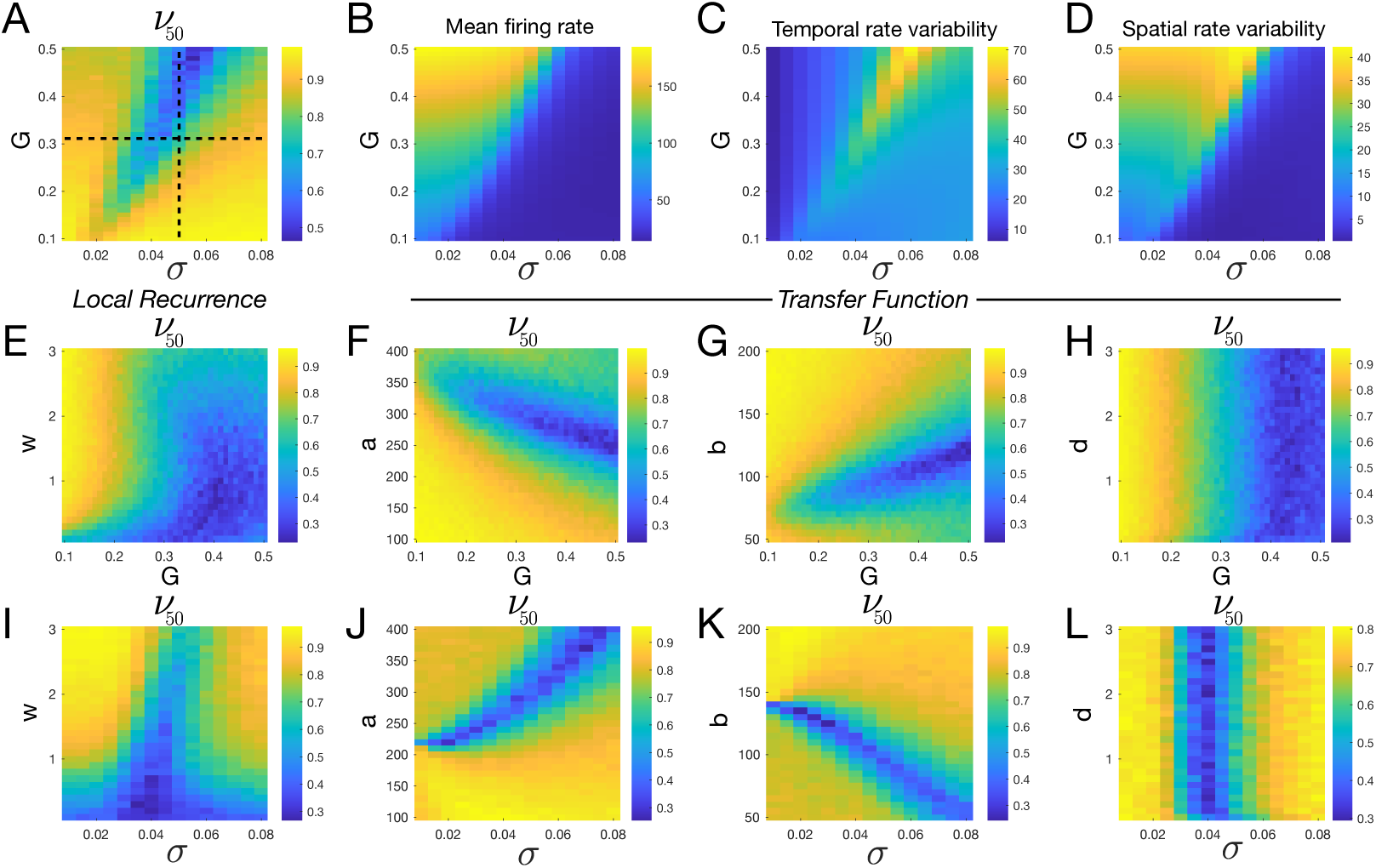
Regimes of faster or slower FCD. (A) Global coupling parameter (*G*) that scales cortical excitability and noise amplitude *σ* modulate FC speed (heat map indicates median speed *ν*_50_ distribution). A level neither too low, nor too large of baseline drive noise *σ* is necessary to arrive at the blue region corresponding to a regime of slower FC dynamics, suggestive of stochastic resonance. (B-D) The mean firing rate, temporal rate variability (mean of the standard deviation of each ROI’s firing rate) and spatial rate variability (standard deviation of the mean of each ROIs firing rate) as a function of G and *σ*. Comparison with panel A indicates that the slower FCD regime corresponds to a regime of dynamic bistability in the firing rate of cortical regimes, akin to spatially distributed transitions between “up” and “down” states. (E-H) Median FCD speed as a function of local model parameters and G. (I-L) FCD speed as a function of local model parameters and *σ*.

**Figure 3:**
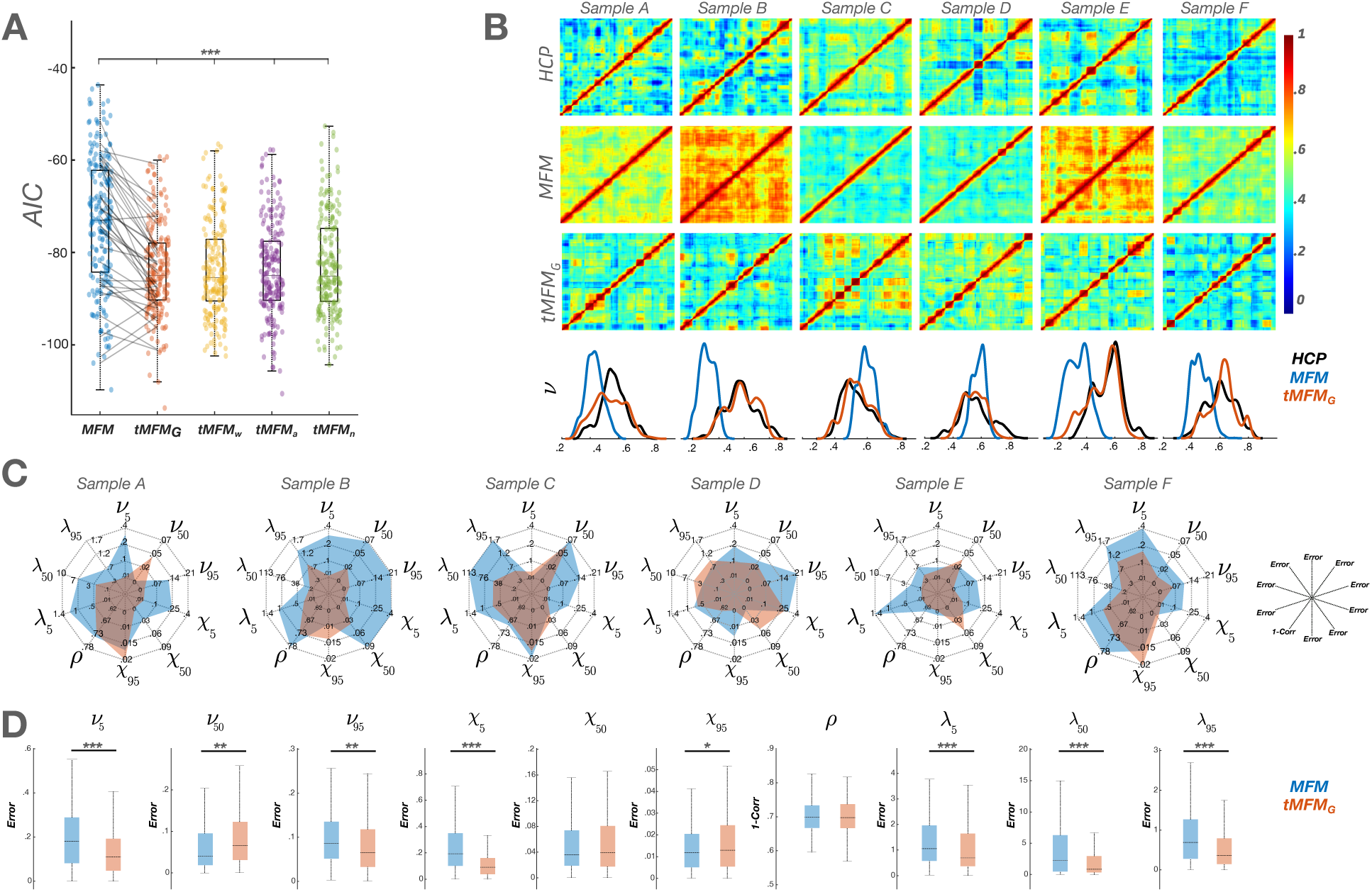
Non-autonomous models outperform autonomous models at explaining detailed FCD features. (A) Comparison of 5 models (1 autonomous and 4 non-autonomous) in their ability to fit a ten-dimensional vector composed of FCD and FC Speed features (5*^th^,* 25*^th^,* 50*^th^,* 75*^th^,* 95*^th^*) percentiles of FCD overall variability *χ* and FCD speed *ν* distributions) derived from 200 HCP recordings. (B) FCD recurrence matrices for six representative sample recordings and the corresponding best fits offered by the MFM and tMFM*_G_* models. In the bottom row, comparison between empirical and model FCD speed distributions for the same five representative subjetcs. (C) Spider plot comparing the ability of the *MFM* (autonomous, blue) and *tMFM_G_* (non-autonomous, red) at capturing the 5*^th^,* 50*^th^* and 95*^th^* percentiles of FCD (*χ*) and FC speed (*ν*) distributions for the 6 samples considered in (B). Additionally, the spider plot indicates how the two models predict match to static FC (*ρ*), computed as correlation distance) and windowed SC-FC correlations (*λ*) that were not used as direct fitting targets. (D) Box-plots indicating the error incurred by each model for both fitting targets (*ν*,*χ*) and prediction targets (*ρ*,*λ*) for the entire dataset consisting of 200 recordings. Results suggest superior quantitative fitting by non-autonomous as compared to autonomous models.

### Predictive power of arousal fluctuations

A good model should not only fit the data it was trained on but also predict independent features that were never part of the fitting process. Adopting this philosophy, we next evaluated how well the non-autonomous, arousal-driven models generalize to other spatiotemporal properties of ongoing brain activity. Specifically, we examined their ability to predict: (i) static functional connectivity (sFC); (ii) the temporal dynamics of structure–function coupling (SC–FC coupling); and (iii) Meta-Connectivity, which captures the covariance among time-varying FC links.

Static FC has historically served as the primary benchmark for whole-brain modelling, with several studies showing that even linear or purely statistical models can capture much of its variance^45,46,47^. Consistent with this, both the autonomous (MFM) and non-autonomous mean-field models (tMFM) reproduced the broad structure of empirical FC, with no significant difference in the correlation *ρ* between simulated and empirical sFC matrices (Fig. 3D).

To detect subtler effects beyond this gross matrix similarity, we further examined how accurately the models captured the range of individual pairwise sFC weights at the single-subject level. For each subject, we extracted the distribution of sFC matrix entries and computed, across recordings, the correlation between empirical and simulated values of the 5*^th^*, 50*^th^*, and 95*^th^* percentiles of these distributions. This analysis allows us to assess how precisely the fitted models capture inter-subject differences in the range of sFC values. Using bootstrap resampling (see *Methods*) with replacement to estimate correlations and confidence intervals, we found that models incorporating arousal-dependent modulation of global excitability markedly improved the prediction of sFC percentiles. This result suggests that abrupt, arousal-induced transitions contribute substantially to shaping mean functional connectivity patterns (see Fig. 4A for bootstrap distributions).

**Figure 4:**
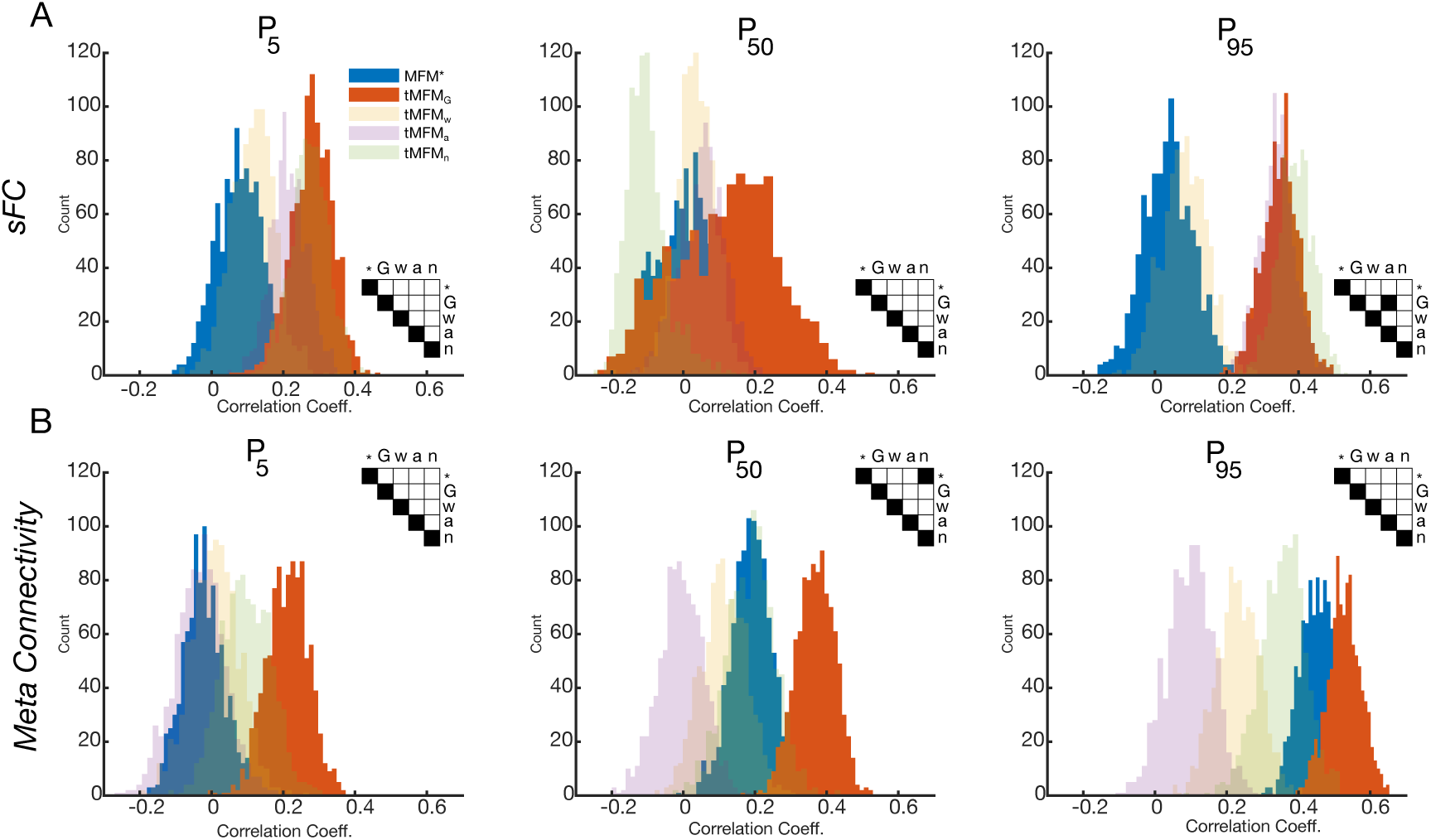
Non-autonomous models outperform autonomous models in novel feature prediction. We assessed how well the different models reproduced empirical FCD features that were not explicitly fitted during model optimisation. We report distributions of boot-strapped correlations between simulated and empirical features, computed for percentiles of the single-subject distributions of (A) static FC matrix entries and (B) Meta-Connectivity entries. Checkered insets indicate significant differences between the correlations achieved by different model types (black = not significant). Non-autonomous models, and in particular the tMFM*_G_*, consistently yield higher correlations.

We next examined the similarity between structural connectivity (SC) and individual frames of the FCD reconfiguration stream. This SC–FC correlation is known to fluctuate over time, reflecting alternations between epochs in which time-resolved FC is more or less constrained by the underlying structural architecture ^48,49^. For this feature, both the autonomous (MFM) and excitation-modulated (tMFM*_G_*) models overestimated the overall level of SC–FC correlation (*λ*), yielding average values of approximately −.004, 0.229, 0.05 and −0.0131, 0.0132, 0.0396, respectively, compared with the empirical mean of −0.142, 0.0082, 0.0305 for the 5*^th^,* 50*^th^,* 95*^th^* percentiles. Despite this offset, the non-autonomous model captured the quantiles of the *λ* distribution with lower percent error, providing a better representation of the distribution tails than of its median (Fig. 3C–D). This improvement suggests that critical bifurcations—enabled by arousal-driven fluctuations—are necessary to reproduce the intermittent decoupling between structure and function observed in human data.

Finally, we turned to Meta-Connectivity^5,11^, a form of edge-based functional connectivity^34^ that quantifies the co-fluctuations of pairwise FC links. Meta-Connectivity (MC) provides a static description of coordinated temporal fluctuations in FC link strengths, in much the same way that classical sFC provides a static description of coordinated temporal fluctuations in regional node activity. The structure of MC, and in particular its modular organisation^34,5^, reveals the existence of sets of links—and hence functional subnetworks—that coherently “pop in” and “out” during the FCD stream. This indicates that FCD is organised not only temporally, but also spatiotemporally. As for sFC, we assessed how well the models reproduced the distribution of MC entries at the single-subject level, using distribution percentiles as prediction targets rather than quantities directly fitted during model optimisation. Analogously to Fig. 4A, we report bootstrapped correlations between the 5*^th^,* 50*^th^, and*95*^th^* percentiles of matched simulated and empirical MC-entry distributions (Fig. 4B). Once again, the tMFM*_G_*—driven by global fluctuations in cortical excitability (*G*)—significantly outperformed both the autonomous and alternative non-autonomous models in accounting for empirical Meta-Connectivity statistics.Specifically, the improvement is most pronounced for the 5*^th^* percentile, corresponding to the lower tail of the MC-entry distribution, which also includes rare negative MC values. This enhanced ability to capture the left tail is particularly important, as the emergence of “viscosity”—that is, the appearance of an increased number of negative MC entries—has been associated with pathological progression in neurodegenerative diseases such as Alzheimer’s disease^11,12^.

Taken together, these results demonstrate that arousal-driven modulation improves not only goodness of fit, but also predictive generalisation across multiple spatiotemporal scales of brain dynamics.

### Neuromodulatory inputs induce working point excursions across critical phase transition boundaries

The superior performance of the arousal-driven model can be traced to how arousal fluctuations navigate the system’s phase space. The mean-field model exhibits distinct regimes demarcated by slower (blue) and faster (yellow) speed of FCD reconfiguration (Fig. 2A). In the arousal-modulated model, fluctuations in the working point allow the system to roam back and forth across these critical boundaries, producing abrupt dynamical transients. This capacity for controlled excursions between regimes appears to underlie the enhanced temporal flexibility captured by the model.

Taken together, the fitting and predictive analyses converge on a simple conclusion: slow, global fluctuations in cortical excitability (captured by the parameter (*G*) provide a parsimonious account of the temporal organization of FCD. To examine this mechanism more directly, we compared the dynamic working points (DWPs) inferred from model fits of autonomous and non-autonomous formulations to empirical resting-state recordings. In the autonomous models, fitting yields a single, time-invariant dynamic working point defined by the optimal values of (*G*) and (*σ*). In contrast, the non-autonomous tMFM models allow selected parameters to fluctuate over time; for each such parameter we estimated a baseline level, a characteristic timescale, and a volatility, which together define both the range of variation and the corresponding occupancy distribution. We focused in particular on the inferred fluctuation range of the global coupling parameter (*G*) in the fitted *tMFM_G_* models.

As shown in Fig. 5A, the dynamic working points (DWPs) inferred for autonomous mean-field models of the five representative subjects in Fig. 3B–C cluster tightly near the critical boundary associated with the model’s “flaring” rate instability, (*G*_+_(*σ*)). This pattern generalizes to the full dataset of 200 recordings (Fig. 5D), where the majority of inferred solutions likewise lie in close proximity to this instability line. Together, these results lend strong support to the longstanding hypothesis that large-scale brain dynamics operate near critical transitions^18,19,20^, at least when described by autonomous whole-brain models.

**Figure 5:**
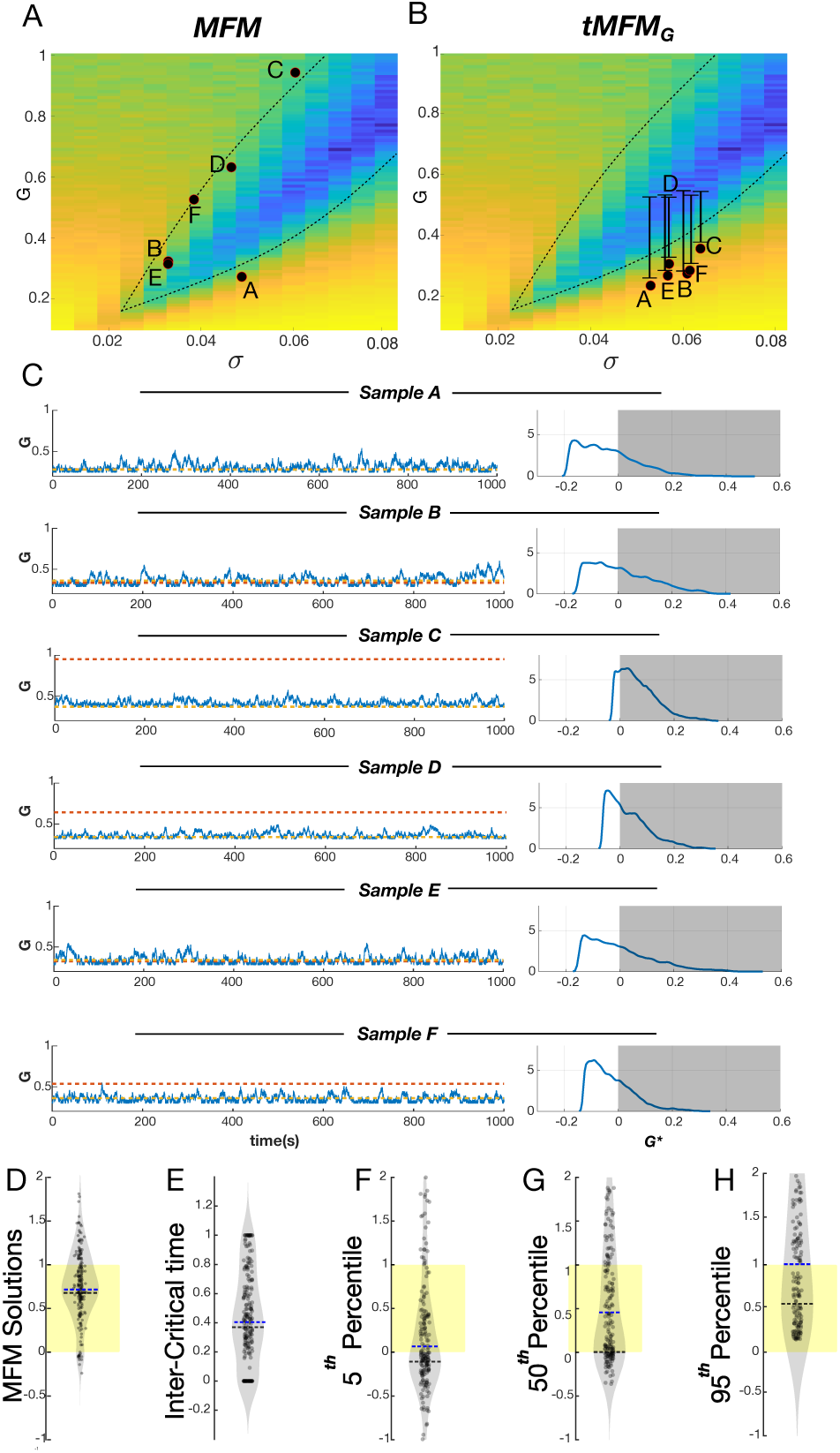
From the critical point to the critical roaming hypothesis. A) Inferred working points for the MFM (autonomous) for six representative subjects; best fit DWPs lie close to the critical flaring rate *G*_+_ instability. As in Fig 2A-D, *G* scales the strength of inter-reginal connections while *σ* is the amplitude of baseline drive noise supplied to each node. Heatmap color indicates the median FCD speed (*< ν*_5_0 *>*). B). For the tMFM*_G_* (non-autonomous model), a range of possible values that a fluctuating *G*∗ can assume is fitted instead. The inferred ranges indicate that the DWP has a baseline level below the critical ignition boundary *G*_−_ and makes transient excursions beyond it. C). (Left) Time series of *G* for the six representative subjects relative to the MFM solution (red) and critical ignition boundary (yellow). (Right) The dwell time distributions of *G*, i.e. cortical excitability *G* scaled relative to the Lower (*G*_−_) and Upper critical boundaries (*G*_+_), with 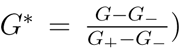. Shaded region lies within the slow speed intercritical range. D) The amount of time spent within the slow regime as a fraction of total area under curve. E) The range of solutions for the autonomous MFM across 200 recordings F-H) The distribution of the 5*^th^*,50*^th^* and 95*^th^* percentile of the normalized *G*^∗^. The average distribution of the percentiles indicates the dynamic range explored by tMFM*_G_*. The dashed blue lines indicate the distribution medians and blacked dashed line the distribution modes. Overall, the fitting of non-automous models suggest that the system spends a substantial part of its time roaming between the ignition and flaring critical transitions.

In contrast, we show in Figure 5B the range between the fitted baseline minimum and maximum values of *G* in the non autonomous tMFM*_G_* for the same five subjects of Fig 5A. In most solutions, the baseline excitability was at values slightly below the ignition critical boundary (Fig 5B,C) (*G*_−_). However, as an effect of parameter fluctuations in time, the system frequently entered the critical zone between the ignition and the flaring lines, spending a substantial fraction of its time within the slow regime (between *G*_−_ and *G*_+_) (Fig 5C, right).

To quantify more precisely and systematically, across the full set of fitted sessions, the range of dynamic working points (DWPs) transiently explored by the system and their relation to the lower ignition and upper flaring critical lines, we introduced a normalized index *G*^∗^ = 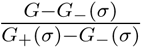. By construction, *G*^∗^ *<* 0 when *G* is subcritical with respect to the ignition point *G*_−_(*σ*) at the fitted value of *σ*; 0 ≤ *G*^∗^ ≤ 1 when the system roams within the slow intercritical regime between the ignition and flaring points *G*_+_(*σ*); and *G*^∗^ *>* 1 when it lies beyond the flaring transition. Fig. 5E shows the fraction of time spent in the slow intercritical regime, which amounts to approximately ∼40%. Figs. 5F–H instead report the quantiles of the single-session distributions of *G*^∗^.

Baseline cortical excitability, tracked by the 5^th^ percentile of the single-session distributin of *G*^∗^, had a median value close to 0, i.e. near the ignition point *G*_−_ (Fig. 5F), while the upper range of excitability, tracked by the 95^th^ percentile of the distribution of *G*^∗^, had a median value close to 1, i.e. near the flaring point *G*_+_ (Fig. 5H). Despite variability across recordings, the tails of the normalized *G* distributions thus consistently indicated subcritical baselines punctuated by frequent excursions beyond the lower ignition critical line. A smaller subset of solutions exhibited crossings of both the lower ignition and upper flaring boundaries, or of the upper boundary alone, reflecting transient transitions into fast, high-excitability regimes (Fig. 5F–H).

Taken together, these results suggest that, in contrast with the critical point hypothesis—according to which the DWP remains close to the flaring rate instability, as implied by the autonomous best-fit models— arousal-driven modulations enable the brain to dynamically explore the entire range between the ignition and flaring lines, intermittently departing from a subcritical baseline below ignition. We refer to this alternative scenario, supported by the non-autonomous model fits, as the *critical roaming hypothesis*.

## Discussion

Spontaneous brain activity is richly structured in time, yet the dynamical principles governing these fluctuations have remained elusive. Here we show that slow, global modulations of cortical excitability—an interpretable proxy for neuromodulatory arousal—provide a parsimonious explanation for the heavy-tailed, intermittently reorganizing structure of functional connectivity dynamics (FCD). By directly contrasting conventional autonomous mean-field models with explicitly non-autonomous, arousal-driven variants, we demonstrate that many hallmark features of ongoing FC—fat-tailed step-length distributions, alternating epochs of persistence and rapid reconfiguration (Fig 2.), and additional out-of-sample signatures such as SC–FC coupling, fit to static FC and Meta-Connectivity (Fig 4.)—emerge only when the model’s dynamic working point (DWP) is allowed to drift over time. In the autonomous case, reconfigurations require the system to hover near intrinsic bifurcations; the non-autonomous model naturally traverses these transitions, greatly expanding the repertoire of accessible large-scale network states (Fig 5.).

A key conceptual advance enabling this result is our explicit focus on the sequential organization of FCD. Building on previous work quantifying FC variability, we introduce FCD Speed as a measure of the stepwise progression of the system through FC space^4,5,11^. Unlike statistics that summarize FCD variability alone ^50,51^, FCD Speed captures the stochastic, random-walk–like structure of FC trajectories and their occasional large “leaps” ^4^.These heavy-tailed excursions turn out to be crucial: improvements in model performance arise almost entirely from the tails of the FCD and FCD-Speed distributions, not their means, which both model families already reproduce well (Fig 3.). This highlights that meaningful dynamical events—those that reshape whole-brain FC—are rare, abrupt, and deeply informative about underlying mechanisms.

The presence of such intermittent, large-scale reconfiguration events naturally raises the question of where the brain operates relative to critical boundaries. Criticality has long been proposed as an organizing principle of neural dynamics, yet its functional role remains debated. Emerging evidence suggests that the brain does not maintain a fixed proximity to a critical point; rather, it approaches or retreats from criticality depending on behavioral and cognitive demands. Simple tasks may even be hindered by critical dynamics, whereas demanding tasks benefit from them^52^, implying that the “distance to criticality” is itself a tunable control variable. Arousal and vigilance are known to shift this distance^53^, and resting-state fMRI—particularly in eyes-closed conditions—captures drifting mixtures of wakefulness and light sleep^54^. This body of work suggests that the brain’s DWP is inherently nonstationary, wandering across a wide swath of state space, such that small fluctuations in arousal can produce disproportionately large changes in functional connectivity. This may explain why linear models approximate static FC well ^45,47,46^ (see Supplementary) but systematically fail to capture the temporal richness of time-resolved FC.

Our fitted simulations reveal how such wandering arises: slow fluctuations in a global excitability-like parameter (captured by the scaling term (*G*)) repeatedly move the system toward and away from rate-instability transitions (Fig 5.). Although several model parameters can, in principle, induce critical transitions, perturbation analyses consistently identified (*G*) as the dominant determinant of FCD temporal structure. Biologically, this resonates with the diffuse ascending projections of major neuromodulatory systems, which act as global gain controllers and are well positioned to modulate large-scale excitability^25,24,55^. Nonetheless, neuro-modulatory influences are not isolated control knobs but coordinated, low-dimensional patterns of receptor- and state-dependent action. The strong explanatory power of (*G*) therefore likely reflects a projection of this low-dimensional biological manifold onto model space, rather than the dominance of any single biophysical mechanism. Incorporating realistic spatial gradients of neuromodulatory receptor densities—known to be systematic across cortex^56,38^—may further enhance the model’s ability to capture fine-grained spatial structure in Meta-Connectivity, which we considered here only in distributional form.

Several additional observations warrant consideration. While the predominant dynamical motif across participants consisted of fast-regime trajectories interrupted by intermittent excursions into the slow regime, we also observed substantial inter-individual heterogeneity. In a subset of individuals, trajectories originated within the slow regime and periodically crossed rate-instability boundaries, suggesting that different brains may occupy systematically distinct positions along an arousal–excitability manifold. Such alternative dynamical pathways may reflect stable differences in baseline arousal, neuromodulatory responsiveness, or cognitive style, and may ultimately map onto meaningful behavioral, age-related, or clinical phenotypes^57,58,59,60^. Converging evidence from neuropathology, imaging, and clinical studies indicates that neurodegenerative disorders—most notably Alzheimer’s disease (AD)—are characterized by early and progressive disruption of ascending neuromodulatory systems. Degeneration of basal forebrain cholinergic nuclei is a hallmark of AD and underlies the long-standing cholinergic hypothesis, with loss of cholinergic neurons and cortical innervation closely tracking cognitive decline ^61,62^. Similarly, the noradrenergic locus coeruleus shows pronounced vulnerability, often degenerating prior to overt cortical pathology and clinical symptoms^63^. Because these systems exert diffuse control over local cortical parameters, their degeneration is expected to constrain the brain’s ability to dynamically traverse critical regimes of network dynamics. Within the framework developed here, such neuromodulatory loss would restrict excursions across dynamical phase boundaries, narrowing the accessible repertoire of functional network states. This may provide a parsimonious mechanistic account of the reduced dynamical fluidity^64^, impaired functional connectivity dynamics^11^, and diminished metastability^65^ consistently reported in AD, and suggests that neuromodulatory decline contributes to cognitive impairment by producing a fundamental impoverishment of large-scale brain dynamics.

We offer potential customizations of the pipeline proposed here. First, although our implementation relied on the bistable Wong–Wang neural mass—a widely used model for large-scale fMRI dynamics^39,15,38^—the broader framework we introduce is not tied to this specific formulation. Oscillatory mechanisms, which are central to EEG, MEG, and LFP signals, could be readily incorporated using Stuart–Landau oscillators or next-generation neural mass models such as the Montbrío mean-field reduction^66,67^. Extending the framework into these oscillatory regimes may offer a principled route for unifying fast electrophysiological rhythms with slow arousal fluctuations^28,68^. Second, we adopted a genetic algorithm for model fitting because it robustly accommodates noisy, computationally expensive simulations and multi-objective optimization without requiring gradient information^44^. This makes it particularly well suited to whole-brain models, whose parameter spaces are often highly non-linear and characterized by sharp dynamical transitions. By treating the model as a black box, this approach avoids explicit assumptions about likelihood functions or the need for specialized training datasets. Recent work has explored simulation-based inference^69,70^ for parameter estimation in whole-brain models, and systematic comparisons of these complementary approaches—in terms of performance, scalability, and interpretability—represent an important direction for future research.

Finally, our findings also carry important implications for the fMRI community, which has long debated the extent to which resting-state FCD is shaped—or contaminated—by arousal-related fluctuations^26,71^. The present framework offers a principled, dynamical-systems–based method for quantifying these contributions. In particular, it provides a mechanistic counterpart to the distinction proposed by Laumann and colleagues^26^, who separated neural contributions to FCD into components related to spontaneous cognition versus arousal. Within our modeling scheme, this dichotomy maps naturally onto autonomous versus non-autonomous dynamics: the former capturing intrinsic, cognition-related fluctuations, and the latter reflecting slow, state-dependent modulations driven by arousal systems.

Together, our findings offer a unified framework in which neuromodulation shapes large-scale brain dynamics by continuously steering the system across inter-critical regions of state space. Rather than treating resting-state activity as noise around a fixed operating point, our results support a view of the brain as an adaptive dynamical system whose working point fluidly evolves on slow timescales, enabling it to flexibly explore, sample, and reorganize its functional connectivity landscape. This perspective provides a foundation for mechanistically linking arousal, critical dynamics, and spontaneous cognition—and for understanding how their disruption contributes to aging, neuropsychiatric conditions, and altered states of consciousness.

## Methods

### Dataset

The dataset analyzed here corresponds to the same subset of the Human Connectome Project (HCP) used in prior work on test–retest reliability^72^. It includes resting-state fMRI recordings from 100 healthy adults collected by the HCP WU–Minn Consortium. Each participant completed two eyes-open resting scans on separate days while fixating a central cross (200 recordings in all). Functional data were acquired on a 3T Siemens Connectome Skyra using a multiband gradient-echo EPI sequence (2-mm isotropic voxels; TR, 720 ms; TE, 33.1 ms; multiband factor, 8), yielding 1,200 volumes per run (14 min 24 s). High-resolution T1- and T2-weighted images (0.7-mm isotropic) accompanied each session. For all analyses, we used the version of this dataset parcellated into 89 anatomical regions using the AAL atlas.

### FC, FCD and FCD Speed

For both empirical and simulated BOLD data, FCD was computed using a standard sliding-window approach. Each time series was segmented into overlapping windows of 60 s, advanced in 2-s steps (58-s overlap), following^15^. Functional connectivity (FC) was estimated within each window, and pairwise correlations between the upper-triangular elements of all windowed FC matrices were assembled into the FCD matrix:

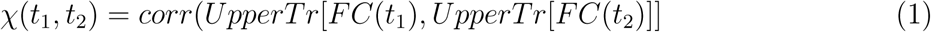

FCD speed was quantified as the rate of change between FCs derived from successive non-overlapping windows:

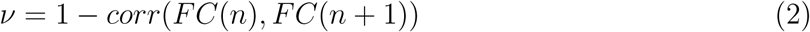

To increase sampling density for relatively short recordings, speeds were computed using window lengths between 55*s* and 65*s* and then pooled. Static functional connectivity (sFC,*ρ*) was computed as the Pearson correlation between the time series of all pairs of ROIs over the entire recording. Time-resolved SC–FC coupling (*λ*) was obtained by correlating each instantaneous FC frame with the structural connectivity matrix. The resulting distribution of (*λ*) values was summarized by its percentiles for subsequent comparison. All analyses were performed using the MATLAB-based *dfcWalk* toolbox using the *TS2dFCstream*, *dFCstream2dFC* and *TS2FC* functions^6^.

### Meta-Connectivity

The idea of functional connectivity can be extended to characterize the dynamic co-fluctuation across all pair-wise links in a network ^6^. Accordingly each inter-regional link is treated as a distinct unit and a correlation matrix is estimated across all *N* (*N* − 1) links (for N ROIs), excluding self-connections, to capture how the fluctuations of one connection co-vary with others over time. This Meta-Connectivity matrix provides a higher-order description of network dynamics, revealing patterns of co-fluctuation between connections in the same way that conventional FC describes correlations between nodes. More formally the Meta-Connectivty between links *ij* and *kl* is given as:

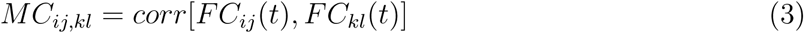

Meta-Connecitivity is computed using the *dFCstream2MC* function of the *dFCWalk* tool-box^6^. In order for robust statistical comparison of static FC fit and Meta-connectivity we devised a bootstrap approach. The 5*^th^*, 50*^th^* and 95*^th^* percentiles of the sFC and Meta-Connectvity distributions were estimated from the HCP data and model solutions for all 200 recordings. A set of 200 random integers ranging between 1 to 200 was generated (allowing for repetition). Pearson Correlation was computed between empirical and model outputs corresponding to each random draw for the sFC and Meta-Connectivity features. This process was repeated 1000 times to obtain a distribution of correlation coefficients for each feature.

### Neural mass modelling

Each ROI is modeled using a mean-field formulation:

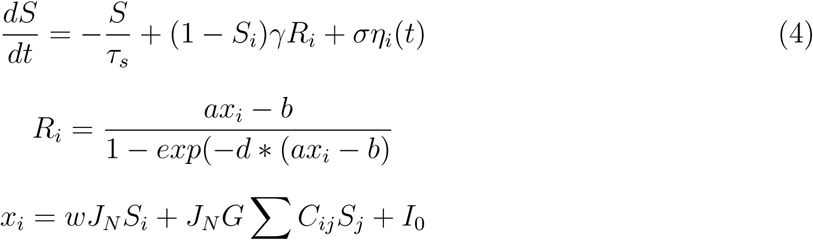

Here, *S_i_* denotes the NMDA synaptic gating variable for region *i*. Total input *x_i_* combines 1. local recurrent coupling scaled by *w*, 2. long-range inputs weighted by the structural connectivity matrix *C_ij_* and globally amplified by the coupling parameter *G*, and 3. a constant external current *I*_0_. Uncorrelated noise enters through a Gaussian term *η_i_*(*t*) with amplitude *σ* and is supplied directly to the synaptic gating term. For the autonomous model (MFM) these parameters assume fixed values for each run. For the class of non-autonomous models (*tMFM*) the parameters *G*,*w*,*a*,*σ* were allowed to fluctuate, one at a time. These slow fluctuations were modeled as an Ornstein–Uhlenbeck (OU) process with mean (*µ*), mean-reversion rate (*θ*), and volatility (*σ*):

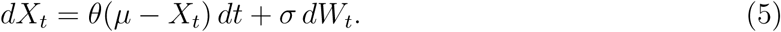

To prevent the process from drifting into physiologically implausible low-arousal states, we imposed a lower bound equal to its mean. Specifically, after each numerical update (Euler–Maruyama), if the process fell below (*µ*), its value was reset to (*µ*):

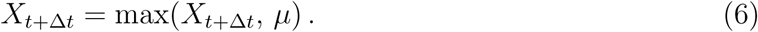

This “truncated” OU formulation preserves the stochastic excursions and autocorrelation structure of the OU process while enforcing a baseline level of arousal consistent with empirical observations. Each simulation was run for (950 s) with a numerical integration step of (1 ms). The initial (50 s) were discarded to eliminate transients. To reduce computational load, BOLD signals were derived by low-pass filtering the firing-rate time series below (0.2 Hz). The validity of this approximation was confirmed by comparison with a standard hemodynamic response function (see Supplementary Materials). Finally, our simulations relied on a publicly available structural connectivity matrix derived from the HCP cohort and parcellated using the AAL atlas https://github.com/juanitacabral/NetworkModel_Toolbox.

### Model Fitting

Model parameters for both the autonomous MFM and the time-varying tMFM were estimated by fitting simulated dynamics to empirical functional connectivity dynamics (FCD). For each model, the fitting targets were the empirical FCD and FCD-speed distributions, summarized by their 5th, 25th, 50th, 75th, and 95th percentiles, yielding a 10-dimensional feature vector capturing the full spread of temporal variability. For each BOLD recording (200 in total) the genetic algorithm (GA) optimized model parameters by minimizing the Euclidean distance between this empirical vector and a corresponding vector computed from simulated data. Optimization was performed in MATLAB using the *ga* function with a population size of 50 and a maximum limit of 100 generations. Default GA operators were used, including scattered crossover and Gaussian mutation, and chromosomes within each generation were evaluated in parallel to accelerate fitness computation. A custom output function recorded the best solution and its corresponding error for each generation for each recording. For each parameter, lower and upper bounds were set based on prior parameter sweeps to constrain the solutions within plausible ranges.

## Acknowledgements

This research has been supported by the Fondation Vaincre Alzheimer (project: **Virtual brains to tailor sensory entrainment and boost memory in early Alzheimer**). We also acknowledge funding by the PEPR Digital Health (project **Brain Health Trajectory**, PEPR ANR-22-PESN-0012-BHT)

