## Supplementary for "The critical roaming hypothesis: arousal-driven transitions across critical lines reproduce human functional connectivity dynamics"

Anagh Pathak

Demian Battaglia

*Université de Strasbourg, CNRS, Laboratoire de Neurosciences Cognitives et Adaptatives (LNCA), UMR 7364, Strasbourg, France*

#### Contents

**1 Comparison with linear dynamics** **2**

**2 BOLD extraction using a hemodynamic response function** **2**

**3 Error incurred by each model class** **3**

**4 Defining critical regimes and boundaries** **4**

### 1 Comparison with linear dynamics

To underscore the idea that purely linear models are incapable of capturing higher order temporal statistics we perform parameter sweep for a linear model where the node dynamics is given by the following equation [1, 2, 3, 4]:

$$\frac{dr_j}{dt} = -r_j + G \sum C_{ij} r_j + \sigma \eta(t) \tag{1}$$

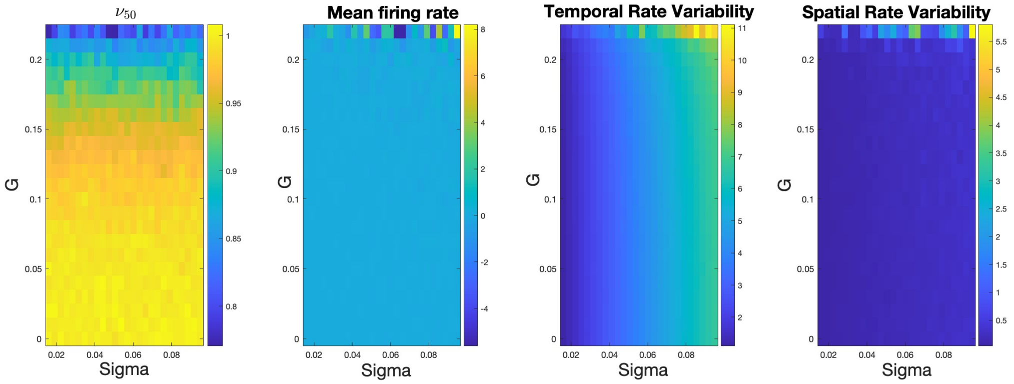


Supplementary Figure 1: Parameter sweep of linear model

Notably, for $G > .22$ the system becomes unstable.

### 2 BOLD extraction using a hemodynamic response function

For our main analyses, we modeled the BOLD signal by applying a simple low-pass filter to the firing-rate time series, which substantially reduces computation time given the complexity of the model simulations and genetic-algorithm fitting. Importantly, we verified separately that using a full HRF convolution yields comparable results (Compare Supplementary Fig 1 with Fig 2A, Main text). To convert simulated neural activity into a BOLD-like signal, we applied a standard hemodynamic response function (HRF) to the neural time series. The HRF was modeled as a canonical double-gamma function (as implemented in SPM [5]), composed of a positive gamma peak at $6s$ followed by a smaller, slower undershoot at $16s$. The HRF was sampled at the native temporal resolution of the neural simulation ($dt = 1ms$) and normalized to unit area.

Neural activity was first convolved with this high-resolution HRF, ensuring that the temporal delay, dispersion, and biphasic shape of the hemodynamic response were accurately captured. After convolution, the resulting high-resolution BOLD estimate was downsampled to the fMRI sampling interval (TR = 1 s) using an anti-aliasing resampling procedure. This approach preserves the temporal fidelity of the neurovascular transformation and avoids aliasing artifacts that would occur if the neural signal were downsampled prior to HRF convolution.

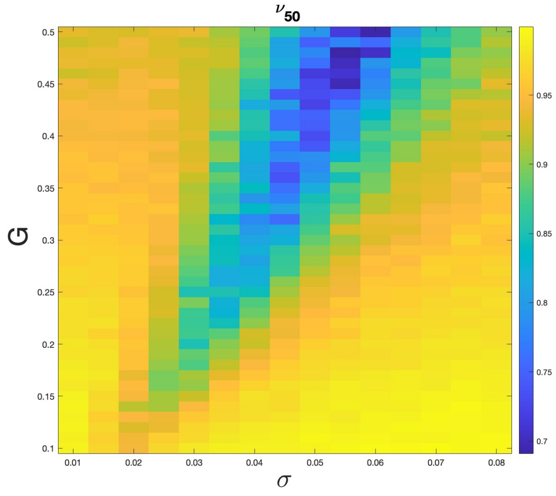


Supplementary Figure 2: Parameter Sweep of $\nu_{50}$ with HRF

### 3 Error incurred by each model class

The figure illustrates the errors produced by each model class over the full dataset of 200 HCP recordings, with each dot representing a single recording. The red line marks the identity line (x = y).

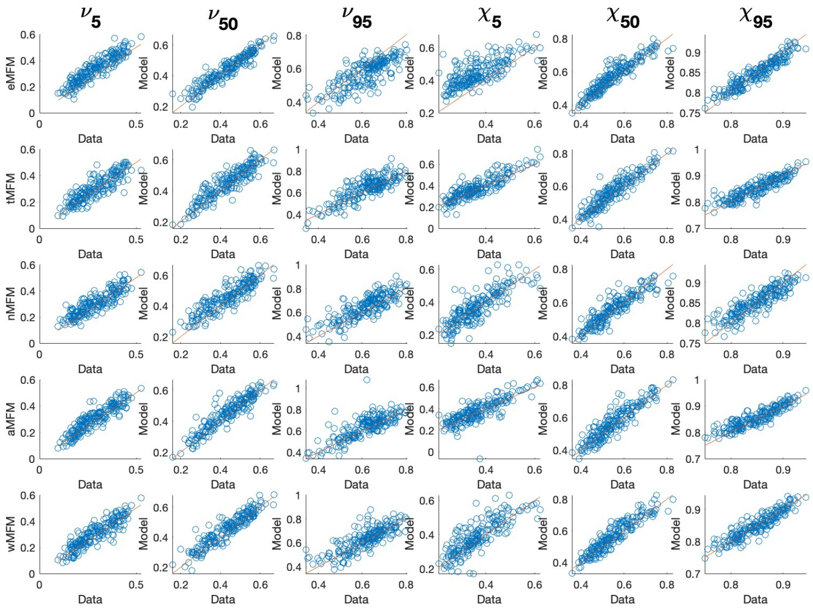


Supplementary Figure 3: Error incurred by each model class

#### 4 Defining critical regimes and boundaries

To delineate critical transition boundaries, we applied a data-driven clustering procedure. For every point in the parameter sweep, we computed a feature vector consisting of the $5^{th}$, $50^{th}$, and $95^{th}$ percentiles of FCD and FCD speed, along with the mean firing rate, temporal rate variability, and spatial rate variability. K-means clustering was then performed on these feature vectors. We evaluated solutions for multiple values of $k$ using the average silhouette score and selected $k = 3$, which yielded the highest score. Boundaries in parameter space were identified using an automated procedure that detected transitions between cluster labels across the sweep. Finally, smooth exponential curves were fitted to these boundary points to define the critical transition lines shown in Fig. 5.

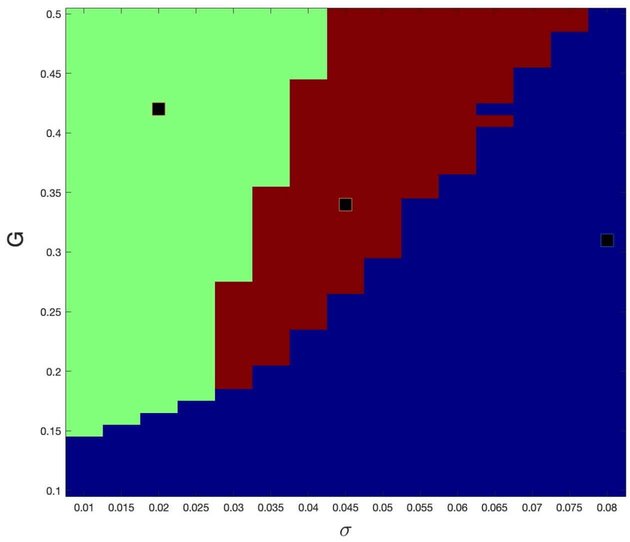


Supplementary Figure 4: k-means clustering in 3 distinct dynamical regimes

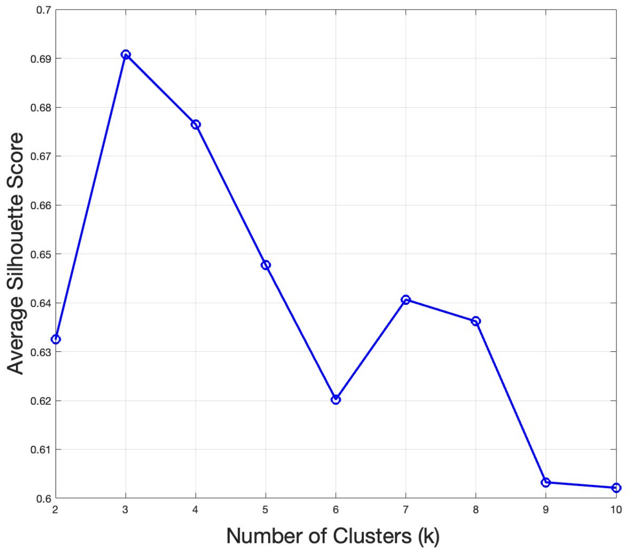


Supplementary Figure 5: Silhouette scores for optimal cluster identification
